# ACDtool: a web-server extending the original Audic-Claverie statistical test to the comparison of large data sets of counts

**DOI:** 10.1101/304568

**Authors:** Jean-Michel Claverie, TA Thi Ngan

**Affiliations:** Structural and Genomic Information Laboratory, Aix-Marseille Université & CNRS UMR7256, Mediterranean Institute of Microbiology (FR3479), Case 934, 163 Avenue de Luminy, Marseille 13288, France

## Abstract

**Motivation:** More than 20 years ago, our laboratory published an original statistical test (referred to as the Audic-Claverie (AC) test in the literature) to identify differentially expressed genes from the pairwise comparison of counts of cognate RNA-seq reads (then called “expressed sequence tags”) determined in different conditions. Despite its antiquity and the publications of more sophisticated software packages, this original article continued to gather more than 200 citations per year, indicating the persistent usefulness of the simple AC test for the community. This prompted us to propose a fully revamped version of the AC test with a user interface adapted to the diverse and much larger datasets produced by contemporary omics techniques.

**Results:** We implemented ACDtool as an interactive, freely accessible web service proposing 3 types of analyses: 1) the pairwise comparison of individual counts, 2) pairwise comparisons of arbitrary large lists of counts, 3) the all-at-once pairwise comparisons of multiple datasets. Statistical computations are implemented using standard R functions and mathematically reformulated as to accommodate all practical ranges of count values. ACDtool can thus analyze datasets from transcriptomic, proteomic, metagenomics, barcoding, ChlP'seq, population genetics, etc, using the same mathematical approach. ACDtool is particularly well suited for comparisons of large datasets without replicates.

**Availability:** ACDtool is at URL: www.igs.cnrs-mrs.fr/acdtool/

**Supplementary information:** none.

## 1 Introduction

Sequence-based approaches started to supersede micro-array hybridization-based platforms for the measurement of gene expression following the introduction of the concept of “expressed sequence tags” (Adams *et al*., 1993). This trend was amplified by the “Serial analysis of gene expression” (SAGE) approach (Velculescu *et al*., 1995; Velculescu *et al*. 1997) that provided an increased output and decreased cost over the sequencing of regular cDNA libraries. At this point, the nature of the raw data collected to quantify the level of gene expression changed from fluorescence intensities to numbers (i.e. counts) of gene-specific sequence tags. Accordingly, new bioinformatic methods had to be introduced to help biologists interpret the data from “digital” gene expression profiling experiments. Our laboratory was among the first to propose a relatively rigorous statistical framework to compare digital expression profiles obtained from two different samples, point out the genes most likely to be differentially expressed, and study the influence of random fluctuations and sampling size on the reliability of these inferences (Audic and Claverie, 1997). This approach, specifically published in the context of investigating differential expression, is in fact applicable to the analysis of counts of any type of event rare enough to be adequately modeled by a Poisson distribution. As the sequence tags approaches became increasingly popular (becoming known as “RNA-seq” following the advent of next generation sequencing systems), more specific bioinformatic packages have been developed reviewed in Hunag *et al*., 2015). Among the most cited are Limma (Ritchie *et al*., 2015), DESeq (Anders *et al*., 2013; Love *et al*. 2014), or EdgeR (Robinson *et al*. 2010; Anders *et al*., 2013). More recently, new bioinformatics packages have been proposed to specifically handle single-cell RNA sequencing data (Kharchenko*et al*., 2014; Finak*et al*., 2015; Sengupta*et al*., 2016; Li and Li, 2018). All the above RNA-seq analysis tools are provided as R/Bioconductor packages the implementation of which requires in-house bioinformatics expertise. To alleviate this requirement, a few tools are now starting to be proposed as web-service to the end-users (Zhang *et al*., 2017; Zhu *et al*., 2017). Surprisingly, despite the lack of follow up, our initial paper (Audic and Claverie, 1997) continued to receive sizable number citations over the years with a large increase since 2012. The continuous usage (hence usefulness) of this statistical test now commonly referred to as the “Audic-Claverie test” (e.g. Bortoluzzi *et al*., 2005; Metta *et al*., 2006; Kim *et al*., 2008; Tino, 2009; Wong *et al*., 2013) prompted us to revisit its mathematical formulation and adapt its numerical calculation to the much larger datasets and count values generated today. We then implemented this modernized R-library-based version of the test as an interactive web-service targeted to biologist end users, allowing three types of analyzes: 1) the basic pairwise comparison of individual counts, 2) the pairwise comparisons of arbitrary large lists of counts, 3) the all-at-once pairwise comparisons of multiple datasets. ACDtool is proposed as an easy-to-use generic tool capable of processing the immense variety of modern omics techniques (RNA-seq, metagenomics, barcoding, proteomics, population genetics, etc) generating very large (albeit often very sparse) count datasets. Given the straightforward and general mathematical principles on which the Audic-Claverie (AC) test was based, ACDtool is not intended to compete with the multiple specialized packages accompanying each of the above techniques. However, the ACDtool web service might remain quite useful to picture the global trends emerging from a given data sets (especially in absence of replicate), and decide whether the amount of information they contain justify the much larger investment required by specialized bioinformatic approaches.

## 2 Methods

We originally introduced our statistical test in the sole context of detecting differentially expressed genes by comparing their cognate tag counts obtained from two different sampling experiments (Audic and Claverie, 1995). ACDtool now extends its application to any sampling protocol where a large number of distinct and independent events (organisms, objects, labels, etc) are detected and counted, each of them representing a small fraction of the total counts. We then made the reasonable assumption that a Poisson distribution is underlying the counts of each of these individual events.

If we perform two sampling experiments, a given event will be counted x times in the first experiment and y times in the second. Audic and Claverie (1995) established that the probability that these counts were generated from the same but unknown Poisson distribution is given by:

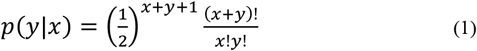

In the general case, where the total numbers of counted events differs in the first (N1) and second (N2) sample, the probability that the counts x and y are generated from samples containing an identical proportion of the corresponding event is given by:

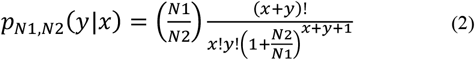

Thus, under the null hypothesis that the tag counts are generated from Poisson distributions with equal means (or proportional to the respective sample sizes), Equation (2) can be used for principled Bayesian inferences, construction of confidence intervals, and statistical testing (Tino, 2009). In the latter case, a cumulative form of Equation (2) (e.g. summing up all the terms in the range [y, 0] if y/N2 <x/N_1_) will be used to compute the p-value.

However, a plain implementation of such simple calculation scheme becomes problematic when applied to the huge range of x and y values encountered in modern omic experiment (RNA-seq, metagenomic, barcoding, etc.).

### 2.1 Unveiling a link with the negative binomial distribution

A significant improvement of our original implementation of the test came after we noticed an unexpected relationship between Equation (2) and the classical negative binomial distribution (NB). Following a little bit of algebra (documented at URL: www.igs.cnrs-mrs.fr/acdtool/) one realizes that

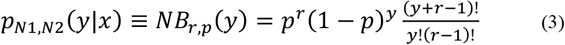

with 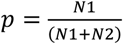 and *r* - 1 = *x*

where *NB_r,p_(y)* is the probability of observing y failures before obtaining r =(x + 1) successes, each one of them with a probability p.

This result calls for two remarks. First, the sampling scheme corresponding to the negative binomial in Equation (3) bears no relationship with the experimental setting at the origin of Equation (2). However, the equivalence between the two expressions nicely establishes a formal link between our Poisson-based initial Bayesian model and the use of the negative binomial distribution arbitrarily assumed for RNA-seq data in subsequent, more specialized, analysis tools (Anders and Huber, 2010; Robinson *et al*., 2010; Di, 2015).

### 2.2 Cumulative form of the negative binomial distribution

An important consequence of the above is that the cumulative form of Equation (2) can be computed using its identity to the negative binomial distribution (Equation (3)) according to:

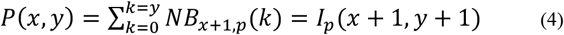

Where I_p_ denotes the incomplete regularized beta function (with valuesin [0-1]), with 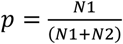

This function is directly available in the R package.

By symmetry, the above formula, valid for 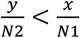, is replaced by

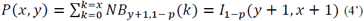

when 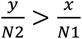.

### 2.3 Introducing a distance

In addition to the calculation of a p-value from a pair of counts associated to the same event (organism, object, label, etc), the ACDtool web service now implements the possibility of comparing whole datasets at once. For this, we introduced a Shannon-inspired measure of distance involving the logarithm of the p-values as computed above, normalized by the relative contribution of each event to the total count. Each event contributes the distance:

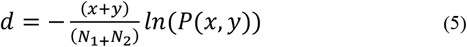

to the global distance D, computed for all K events as:

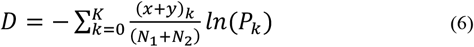

### 2.4 Computing p-values using the hypergeometric function

To avoid potential numerical problems arising from very small p-values, and/or the large range of x, y values, a peculiar attention was devoted to the calculation of Equation (4), (4') and (5). Using standard R functions, a robust implementation was found to use the following formula (Boik and Robinson-Cox, 1999):

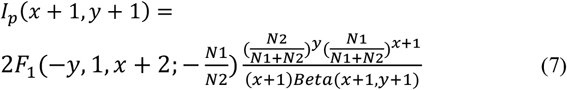

Where 2 *F*_1_(*a, b; c; z*) is the ordinary hypergeometric function of z, with parameters a, b and c. The standard hypergeometric2F1 R function is called with the method flag set as “Laplace”.

## 3 Results

The new ACDtool web service is using the AC statistics in three different contexts. Those different usages are extensively described in the online documentation (accessed via the button tags “document” and “example”). They correspond to the three tools displayed on the home page (URL: www.igs.cnrs-mrs.fr/acdtool/). Their purposes are presented below.

### 3.1 Tool 1: comparing a pair of counts

The input screen of Tool1 requests a pair of counts characterizing the same event (organism, object, label, etc) and the total counts of the two samples from which they were drawn. Each count must be small enough before the corresponding total counts (e.g. smaller than 5%) to justify our assumption that each event is drawn according to its own Poisson distribution. Tool1 then return the probability (and its logarithm) that the compared samples contain the same proportion of the event (Equation (7)). At variance with other inference methods, all reasonable integer values (e.g. in [0, 10^6^]) return a valid result. Tool1 is also helpful at the stage of experimental design to determine which combination of counts and sample sizes are required to diagnose differences reaching a given threshold of statistical significance.

### 3.2 Tool 2: comparing lists of paired counts

This new tool performs the pairwise comparison of two lists of counts associated to the same set of events (organisms, objects, labels, etc) drawn from two samples, and determine which events exhibit the most significant differences. The minimal input format is a count table: the first column lists the different labels and subsequent columns list the counts associated to these labels. The first line of the table displays the heading of each column: label, then sample names. If needed, accessory columns can be embedded to display non-numerical attributes associated to each label. ACDtool2 is expecting a tab-delimited file such as that produced by the Excel spreadsheet (“save as” tab-delimited text, .txt). Such format allows a convenient back-and-forth between Excel editing and ACDtool analyses.

The input screen of Tool2 requests 1) the count table file name, 2) the headings of the two columns of counts to be compared, 3) optionally, the heading(s) of the lists of attributes one wish to add to the output.

The output is an interactive display of the events ranked by increasing p-values (Equation (7)), the sense of (proportional) variation, the original counts, the corresponding normalized distances (Equation (5)), and the selected accessory attributes, if any. This output can be saved as a tab-delimited file (.txt) compatible with the Excel spreadsheet.

### 3.3 Tool 3: evaluating the pairwise distances of multiple datasets

This new tool performs the complete set of pairwise comparisons of multiple lists of counts (associated to the same set of events) all at once, delivering an interactive heatmap of their relative distances (Equation (6)). The corresponding distance matrix can be saved as a tab-delimited file (.txt), for further use such as an input for various clustering algorithms. The input screen of Tool3 solely requests a count table file name. At variance with Tool2, Tool3 will processed all columns as counts, except for the first one. The accessory columns tolerated by Tool2 must then be removed (e.g. using the Excel spreadsheet) from the input file.

**Fig.2.**
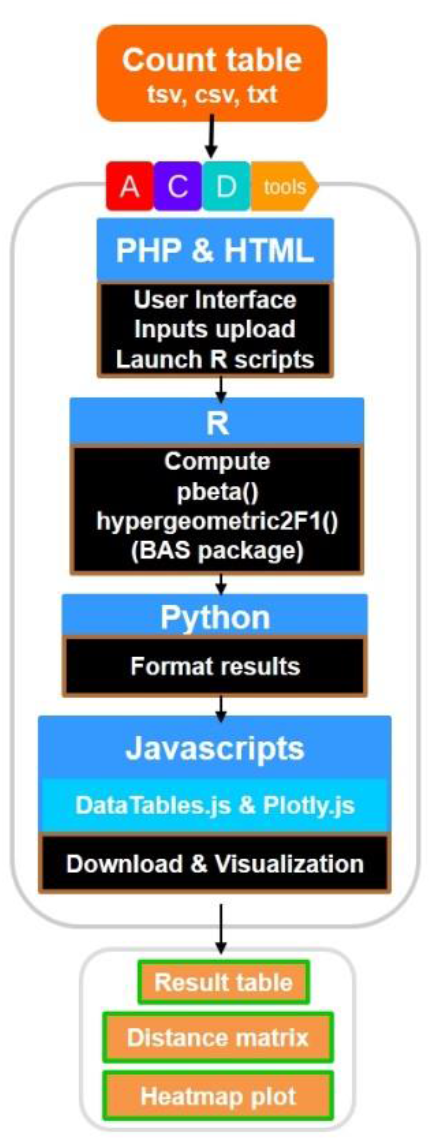
ACDtool implementation.

Tool3 and Tool2 are often complementary. First, Tool3 will be used to reveal the overall similarity/discrepancy between a large number of sampling experiments. Tool2 will then be used to identify which of the events (organisms, objets, labels, etc) are the most discrepant between them.

## Acknowledgements

We thanks Dr. Chantal Abergel for suggesting important improvements to the webserver user interface.

